# Acute Myeloid Leukemia with deletion 5q is an epigenetically distinct subgroup defined by heterozygous loss of *KDM3B*

**DOI:** 10.1101/2024.11.13.623380

**Authors:** Katherine Kelly, Linda Welte, Etienne Sollier, Anna Riedel, Fiona Brown-Burke, Michael Scherer, Harold N. Keer, Mohammad Azab, Ekaterina Jahn, Hartmut Döhner, Konstanze Döhner, Pavlo Lutsik, Christoph Plass

**Affiliations:** Division of Cancer Epigenomics, German Cancer Research Center (DKFZ), Im Neuenheimer Feld 280, 69120 Heidelberg, Germany; Ruprecht Karl University of Heidelberg, Heidelberg, Germany; Department of Internal Medicine III, University Hospital Ulm, Ulm, Germany; Astex Pharmaceuticals, Inc., Pleasanton, CA, USA; Department of Oncology, KU Leuven, Leuven, Belgium

## Abstract

Acute myeloid leukemia (AML) is a hematological malignancy characterized by a block in differentiation and accelerated proliferation of myeloid progenitor cells. Genes encoding for epigenetic regulators are among the most frequent targets for mutations and structural variations in AML, giving rise to profound epigenetic heterogeneity between and within tumors. Deletions of chromosome 5q [del(5q)] are among the most common copy number alterations in AML and are associated with extremely poor clinical outcome and therapy resistance, however the mechanisms linking del(5q) to leukemic progression are not understood. Analyzing DNA methylation profiles from 477 elderly AML patients using DNA methylome deconvolution, we discovered that del(5q) AML is an epigenetically distinct subgroup characterized by a signature of DNA hypermethylation, which we propose may be linked to dysregulation of H3K9me1/2 and overexpression of the leukemic stem cell marker, *DNMT3B*. Interrogation of the minimally deleted 5q region highlighted the H3K9me1/2 demethylase KDM3B as a likely target for haploinsufficiency in this subgroup. Our data suggest that del(5q) AML should be reconsidered as an epigenetically dysregulated subgroup, driven by heterozygous loss of *KDM3B*, and that the resulting imbalance of H3K9me1/2 may contribute to the progression of these aggressive leukemias.

## Introduction

Acute myeloid leukemia (AML) is an aggressive hematological cancer characterized by the accumulation of abnormally differentiated myeloid progenitor cells in the bone marrow and blood. Despite usually having a low mutational burden and few cytogenetic abnormalities compared to other cancer types, epigenetic dysregulation is recognised as a defining feature of this disease ^1^. Genes encoding epigenetic enzymes, such as regulators of DNA methylation, histone and chromatin modifications, are among the earliest and most frequent targets for mutations, structural rearrangements, and copy number alterations (CNAs) in AML^2,3,4,5,6,7,8^. These aberrations can disrupt global epigenetic regulation, conferring inter- and intra-tumor variation in DNA methylation and chromatin architecture. Epigenetic alterations in AML may disturb normal hematopoietic differentiation trajectories, and can ultimately give rise to transcriptional, functional, and clinical heterogeneity ^1,9^.

This background has prompted several previous attempts to define epigenetic subgroups in AML by applying clustering methods to bulk DNA methylation data. For example Figueroa *et al.* identified 16 methylation-based AML subgroups using the promoter-centered HpaII tiny fragment enrichment by ligation-mediated PCR (HELP) assay ^10^, which were later redefined to 14 subgroups using Enhanced Reduced Representation Bisulfite Sequencing (ERRBS) ^11^. More recently, Giacopelli *et al.* defined a set of 13 methylation-based subgroups, so-called “epitypes” using HumanMethylationEPIC (EPIC) and HumanMethylation450K (450K) array data from the BEAT and TCGA AML cohorts ^12^. These studies and others have succeeded in linking methylation signatures to mutations in *IDH1/2* ^5^, *DNMT3A*, *NPM1*, and *CEBPA*, as well as inv(16), t(15;17), t(8;21) translocations, overexpression of *MECOM/EVI-1*^13^*, KMT2A/MLL* rearrangements ^14^, and complex karyotype (ckAML)^12^.

CkAML is an aggressive subgroup defined by the presence of at least three unrelated chromosome abnormalities in the absence of other class-defining genetic alterations, which is enriched for deletions in 5q, 7q, and 17p, as well as TP53 mutations ^14,15,16^. Del(5q) is one of the most common CNAs in AML, occurring in 60-80% of ckAML patients and occasionally as a sole abnormality^15^. Del(5q) is an early event in leukemogenesis, evidenced by its typical monoclonality and frequent occurrence in premalignant myelodysplastic syndromes (MDS) which can progress to ckAML ^15^. Earlier studies attempting to pinpoint the target of this deletion have led to the definition of a CDR at 5q31.2 ^17–20^. This is distinguished from the more telomeric locus associated with low-risk MDS or “5q syndrome”, which has been linked to haploinsufficiency of the ribosomal gene RPS14 ^21,22^. Since no genes within the 5q31.2 locus have shown evidence of mutational inactivation of the second allele, the del(5q) phenotype in high-risk AML is also believed to be driven by haploinsufficiency of one or more genes ^17,18,23–25^. Several candidates have been investigated, most notably *CTNNA1* and *EGR1*^17,18,23–25^, yet, attempts so far to delineate the mechanism underlying the 5q deletion have been inconclusive.

Elderly AML patients usually harbor different molecular and cytogenetic alterations compared to younger individuals, including a higher frequency of ckAML^7,26^. Coupled with the fact that shifts in DNA methylation, including that of hematopoietic stem and progenitor cells (HSPCs), is an established hallmark of aging ^27,28^, the epigenetic landscape in elderly AML requires separate investigation.

Recent studies of DNA methylation have led to clinical advancements in certain cancer types, where epigenetic profiling has been employed for diagnostic and prognostic classification, and to guide therapy selection ^29,30^. Similar applications may be within reach in AML, however, one major challenge in such analyses is the fact that methylation data coming from a single patient likely carries a mixture of epigenetic signals derived from non-cancerous cells in the bone marrow as well as different populations of leukemic cells, often with divergent mutational processes impacting the methylome. This heterogeneity makes it difficult to decipher the causes and consequences of epigenetic disturbances in cancer, and may hinder the effective subclassification of patients. We hypothesized that a methylome deconvolution approach may offer advantages over bulk methylome analysis for the molecular classification of heterogeneous tumors. Methylome deconvolution aims to extract cell-type-specific information and other recurring sources of epigenetic variation that might otherwise be obscured within the bulk cancer methylome ^31^. Such a method would allow us to explore inter-patient heterogeneity, as well as to disentangle the epigenetic heterogeneity within each tumor and its microenvironment. With this approach, we aimed to assess the utility of DNA methylation profiles to stratify elderly AML patients into biologically and clinically relevant subgroups, and to improve existing molecular classification and understanding of AML. This analysis led us to identify an epigenetic signature enriched in patients with del(5q), which we suggest is linked to the heterozygous loss of the H3K9me1/2 demethylase KDM3B and overexpression of *DNMT3B*.

## Results

### Methylome deconvolution-based characterization of AML reveals subgroups defined by distinct molecular and cytogenetic features

To characterize the methylation landscape of AML in older individuals, we focused on a cohort of 477 untreated patients [median age 77, range (59–94)] who participated in the ASTRAL-1 clinical trial ^7,32^. We utilized DNA methylation profiles generated from bulk bone marrow and peripheral blood samples using the Illumina Infinium EPIC array. Subsequently, we applied MeDeCom - a reference-free methylome deconvolution method we earlier developed ^31,33^ - to decompose the original bulk methylation matrix into a set of 11 Latent Methylation Components (LMCs). These LMCs represent major sources of epigenetic variation in our dataset, potentially reflecting cell-type-specific signals, clinical features, and molecular phenotypes (see Methods for details). We interpreted and annotated the resulting LMCs (Figure 1A, Figure S1A, B) by comparisons to known cell types/states, and to mutational and cytogenetic features of tumors (Table S2).

**Figure 1.**
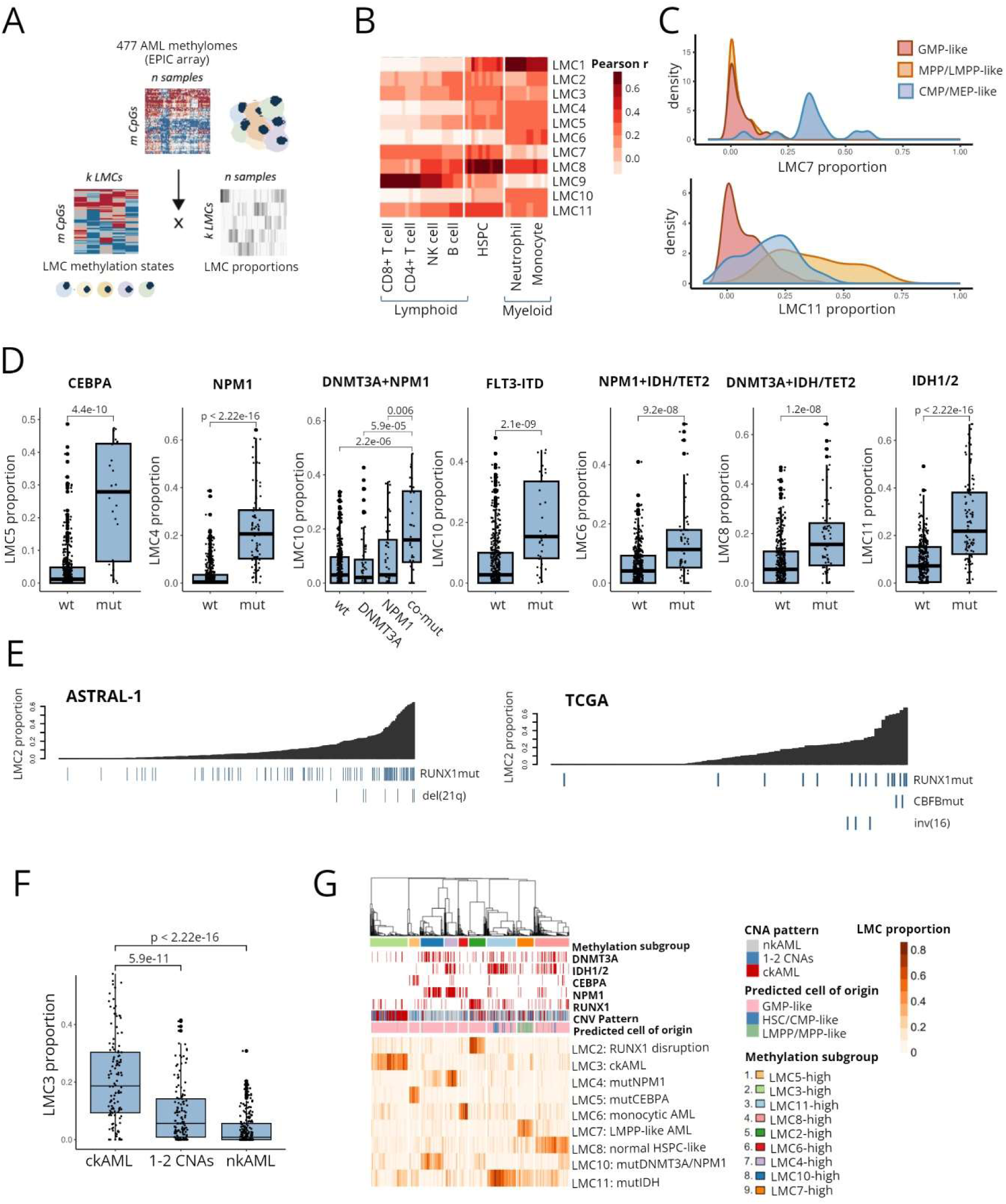
Epigenetic characterization of AML by methylome deconvolution reveals subgroups defined by distinct molecular and cytogenetic features. **A.** Schematic outlining the methylome deconvolution approach and its application to our EPIC array dataset. Methylation data measured from bulk samples is represented as a matrix with n columns (samples) and m rows (CpG sites). MeDeCom decomposes the matrix to derive two further matrices; one of LMCs (k LMCs x m CpGs) and one of LMC proportions (n samples x k LMCs). **B.** Heatmap showing correlation of AML LMCs to normal hematopoietic cell methylomes. Color intensity represents Pearson correlation coefficient. **C.** Density plots comparing LMC proportions in AML methylomes predicted to derive from GMP-like (257), MPP/LMPP-like (47) and CMP/MEP-like (13) cells of origin. **D.** Boxplots showing association of selected LMCs with mutations in epigenetic regulators. Wilcoxon’s p-values shown. **E.** Barplot showing association of LMC2 with RUNX1/CBFB disruption in our own and TCGA cohorts. Patient samples are ordered by increasing LMC2 proportion. **F.** Boxplots comparing LMC3 proportion in relation to patterns of copy number variation. Patients are classified as normal, intermediate or complex karyotype according to the presence of CNAs as detected by Conumee [normal karyotype (nk)AML; no CNAs, intermediate karyotype (ik)AML; 1-2 CNAs, complex karyotype (ck)AML; ≥ 3 CNAs]. Wilcoxon’s p-values shown. **G.** Heatmap of LMC proportions used to define methylation-based subgroups by consensus k-means clustering. Lymphoid-like (LMC9) and neutrophil-like (LMC1) components, and low purity samples (lower tertile InfiniumPurify score) are excluded for clustering.

We first identified LMCs which were derived from non-leukemic cells and differentiation stages by correlating the matrix of LMCs against a range of normal hematopoietic cell methylomes ^34^. We found that LMC1 resembled methylomes of healthy mature myeloid cells (Figure 1B), and was enriched for hypomethylation of neutrophil/macrophage-related gene sets (Table S3). LMC9 correlated with lymphoid cell methylomes (Figure 1B), and with measures of sample purity (Figure S1C) ^35,36^ and was hypomethylated for various immune-related genes (Table S4), while LMC8 closely resembled the methylomes of untransformed HSPCs (Figure 1B).

Since AMLs can also derive from epigenetically distinct HSPC stages ^37,38^, we expected certain LMCs to reflect methylation patterns inherited from their respective ancestral cells. To test this, we estimated the cell of origin of each AML methylome by comparison to various normal HSPC states (see Methods for details). This resulted in three groups: GMP-like, MEP/CMP-like, and MPP/LMPP-like, with the majority of samples closely resembling the GMP state (Figure S1D). We then compared LMC proportions between the resulting groups and found that LMC7 and LMC11 were enriched in the more primitive MEP/CMP-like and MPP/LMPP-like AML, respectively (Figure 1C).

Next, we tested whether certain LMCs were associated with the mutational status of epigenetic regulators or other genetic alterations. We found that LMC4 was enriched in *NPM1*-mutated samples, and LMC5 in AML with *CEBPɑ* mutations (Figure 1D). LMC10 was enriched in samples carrying co-occurring mutations in *DNMT3A* and *NPM1* (Figure 1D), and captured a signature of genome-wide hypomethylation (Figure S1E), as previously described for *DNMT3A*-mutated AML ^39^. The same component was increased in *FLT3-ITD* mutated AML (Figure 1D) and was enriched for hypomethylation at STAT transcription factor binding sites (TFBSs) as previously described for this subgroup (Table S5) ^12^. LMC11 captured a genome-wide hypermethylation signature (Figure S1E) consistent with its enrichment in IDH1/2 mutated AML ^40^. Patients with co-occurring *DNMT3A*+*IDH/TET2* mutations, rather than carrying either global hyper/hypomethylation signatures, were enriched for the normal HSPC-like LMC8 (Figure 1D). This pattern is reminiscent of the “epigenetic antagonism” described by Glass et al., who showed that *IDH1/DNMT3A* dual mutation resulted in a methylation landscape similar to normal CD34+ cell ^11^. LMC6 was associated with *MLL/KMT2A* rearrangements and monocytic M1 FAB classification (Figure S1F), and was enriched in samples with co-occurring *NPM1/TET2* and *NPM1/IDH* mutations. LMC2 was found in patients carrying a range of alterations convergent on the disruption of RUNX1 (Figure 1E). This component was especially high in a subset of ckAML harboring *RUNX1* mutations and/or del(21q), in which the minimally deleted region has been mapped to the *RUNX1* locus ^41^. Additionally, LMC2 was enriched in patients with mutations in the RUNX1 binding partner CBFβ, and in inv(16) AML in which the CBFβ protein is disrupted. Consistent with this, both RUNX1 and CBFβ TFBSs were enriched among LMC2-hypermethylated CpG sites (Table S6). Finally, LMC3 was enriched in patients with ckAML (Figure 1F).

Having linked each LMC to a defining cellular, molecular, or cytogenetic feature, we used LMC proportions to define methylation-based subgroups by applying consensus k-means clustering (Figure S2)^42^. Neutrophil-like LMC1 and lymphoid-like LMC9 were removed from the clustering, since they likely represent confounding by non-leukemic cells. This resulted in nine clusters (see Methods for details) each dominated by a single LMC (Figure 1G, Table S2). In summary, this methylome deconvolution approach allowed us to identify and biologically interpret the principal sources of DNA methylation heterogeneity in elderly AML, and thereby stratify elderly AML patients into biologically meaningful subgroups.

### A DNA hypermethylation signature highlights del(5q) AML as an epigenetically distinct subgroup

Since an in-depth epigenetic characterization of ckAML has not yet been conducted, we decided to explore the origin of the LMC3 methylation signature, which we found to be enriched in ckAML. Specifically, we aimed to investigate whether any specific CNAs might differentiate LMC3-high from LMC3-low ckAML. A comparison of LMC3 proportions against each individual chromosome/arm-level CNA revealed del(5q) as the only subgroup of ckAML in which this signature was significantly enriched (Figure 2A-B). We validated this association using estimated LMC proportions in patients from both TCGA and BEAT-OSU ^12^ AML cohorts (Figure S3A). We showed that patients with isolated del(5q) also carried significantly higher LMC3 proportion than other samples without ckAML (Figure 2B). We verified that this association was independent of TP53 mutation status (Figure S3B), and was not related to other common CNAs, which frequently co-occur with del(5q) (Figure S3C). We used methylome data from sorted AML leukemic stem cells (LSCs) and blasts ^38^ to confirm the presence of LMC3 in both LSC and blast counterparts of del(5q) samples, with higher levels in del(5q) LSCs (Figure S3D). We noted that LMC3 was absent in two del(5q) samples, which we decided to investigate as outliers: one of these samples carried a whole genome duplication (i.e. retaining three copies of 5q), and the other contained a deletion spanning only the distal region and not the 5q commonly deleted region (CDR) (Table S2). These observations suggest that the loss of some entity from the deleted region likely causes this epigenetic signature to arise.

**Figure 2.**
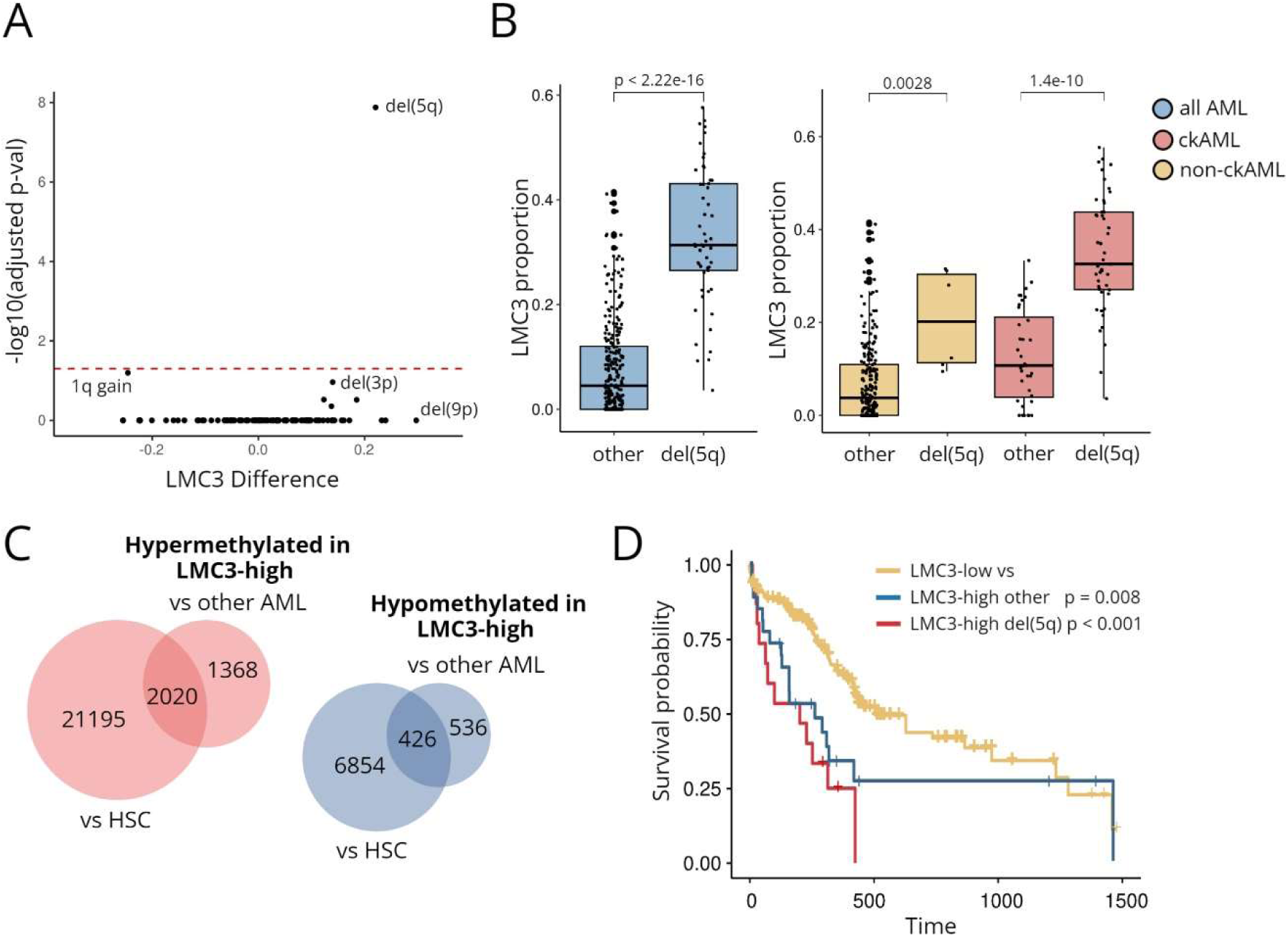
A unique DNA hypermethylation signature distinguishes del(5q) AML from other complex karyotypes. **A.** Volcano plot summarizing the comparison of LMC3 levels against each copy number gain and loss among our ckAML patients. The x-axis depicts the difference in mean LMC3 proportion between patients with and without the specified CNA, and the y-axis shows the -log10 transformed, bonferroni-corrected Wilcoxon p-value for each comparison. The dotted red line indicates an adjusted p-value of 0.05. **B.** Boxplots comparing LMC3 proportion in patients with and without del(5q), among all (left), all non-ckAML (middle) and all ckAML (right) patients from our cohort. Wilcoxon’s p-values shown. **C.** Venn diagram showing the number of overlapping CpG sites detected as hypo/hypermethylated in the LMC3-high AML subgroup by comparison to other AML subgroups and normal hematopoietic stem cells (HSC). **D.** Kaplan Meier survival plot comparing overall survival of patients in the BEAT AML cohort. Patients are separated into LMC3-high and LMC3-low groups based on mean LMC3 proportion, and LMC3-high patients further stratified as del(5q) or otherwise. Pairwise log rank p-values shown.

Given that del(5q) AML has not previously been studied from an epigenetic perspective, and the target of the deletion remains elusive, we hypothesized that LMC3 might shed light on the molecular events linking del(5q) to leukemic progression. To better understand this epigenetic signature, we investigated the CpG sites that were differentially methylated in the LMC3-high subgroup. Taking the overlap of differentially methylated sites from comparison to all other AML subgroups and to normal HSCs, we defined a set of 2020 hypermethylated and 426 hypomethylated CpGs (Figure 2C, Table S7, Table S8). Since hypomethylated CpGs were relatively rare and showed few enrichments of gene ontology terms or TFBSs, we focused our analysis on the hypermethylated sites. LMC3-hypermethylated sites were distributed throughout CpG islands and shores as well as at CpG poor intragenic regions and gene bodies (Figure S4A). They were almost completely unmethylated throughout the normal hematopoietic lineage (Figure S4B). Hypermethylated CpGs were enriched for sites of H3K27me3 and PRC2 binding sites (Figure S4C); a pattern which has been linked to stemness in other cancer types ^43^. Various developmental gene sets were also enriched (Table S9), including an abundance of homeobox genes (Figure S4D), which have been recognized as essential players in normal and malignant hematopoiesis ^44^. We also detected similar developmental processes among the top transcriptional changes which correlated with LMC3 (Table S12).

To clarify the clinical significance of LMC3 beyond the known prognostic indication of del(5q) itself, we took advantage of clinical and drug sensitivity data from the BEAT-AML cohort, which contained a sufficiently large number of LMC3-high patients both bearing and lacking del(5q). Irrespective of del(5q) status, we found that LMC3-high patients had significantly lower overall survival (Figure 2D) and overall reduced drug sensitivity (Figure S4E). The only other LMCs which had prognostic significance were LMC5, which was associated with a favorable outcome, consistent with the enrichment of *CEBPA* mutations, and, to a lesser extent, LMC10 (Figure S5).

In conclusion, we identified a unique hypermethylation signature associated with del(5q) AML, which appears to hold prognostic value independently of del(5q) itself.

### Interrogation of the 5q commonly deleted region implicates KDM3B as a likely haploinsufficiency candidate

A recurrent chromosomal deletion in cancer is typically viewed as the location of a tumor suppressor gene. Attempts to delineate the mechanisms of del(5q) AML have localized several candidate tumor suppressor genes to the CDR at 5q31.2, however there remains little mechanistic evidence linking the loss of any candidate gene to leukemic progression ^17–19^. Finding a methylation signature enriched in del(5q) patients encouraged us to consider a possible epigenetic basis for this aberration. We hypothesized that the loss of an epigenetic regulator encoded by the CDR might explain the observed hypermethylation signature.

To narrow down a CDR in our AML cohort, we analyzed the overlap of deleted segments using copy number profiles estimated from the DNA methylation data from all 79 del(5q) cases ^45^. This resulted in a CDR containing 20 genes, flanked by *MYOT* and *SIL1*, in agreement with previous definitions (Figure 3A, Table S13). Correlation analyses of the resulting CDR genes in two independent datasets (our and TCGA cohorts) identified three candidates whose expression significantly and consistently correlated with LMC3: *KDM3B*, *CTNNA1*, and *ETF1* (Figure S6A, Table S14). We noted that the H3K9me1/2 demethylase *KDM3B* was the only one of the downregulated genes contained within all versions of the CDR described in the literature ^17,20,46,47^, which by its smallest estimate has been narrowed down to an interval containing only *KDM3B, EGR1*, and *REEP2* ^20^ (Figure 3A). Moreover, differential gene expression analyses comparing del(5q) to other AML patients identified *KDM3B* as the most significantly downregulated of all genes within the CDR in both our ckAML (Figure S6B, Table S15) and TCGA cohorts (Figure 3 A-B, Table S16). Using the Ng *et al.* dataset ^48^, we confirmed that *KDM3B* expression is downregulated in both LSCs and blasts from del(5q) samples (Figure S6C). Using two published proteomics datasets from AML patients ^49,50^ as well as data from AML cell lines ^51^, we consistently observed a significant reduction in KDM3B protein expression in the del(5q) subgroup, which was not consistently observed for other notable CDR candidates, CTNNA1, EGR1 or ETF1/eRF1 (Figure 3C & Figure S6D-E). From pan-cancer and pan-tissue analyses of TCGA and GTEx data, we observed that *KDM3B* expression is higher in AML than in any other tumor or normal tissue type (Figure S6F), suggesting that it may be an important gene in this disease context.

**Figure 3.**
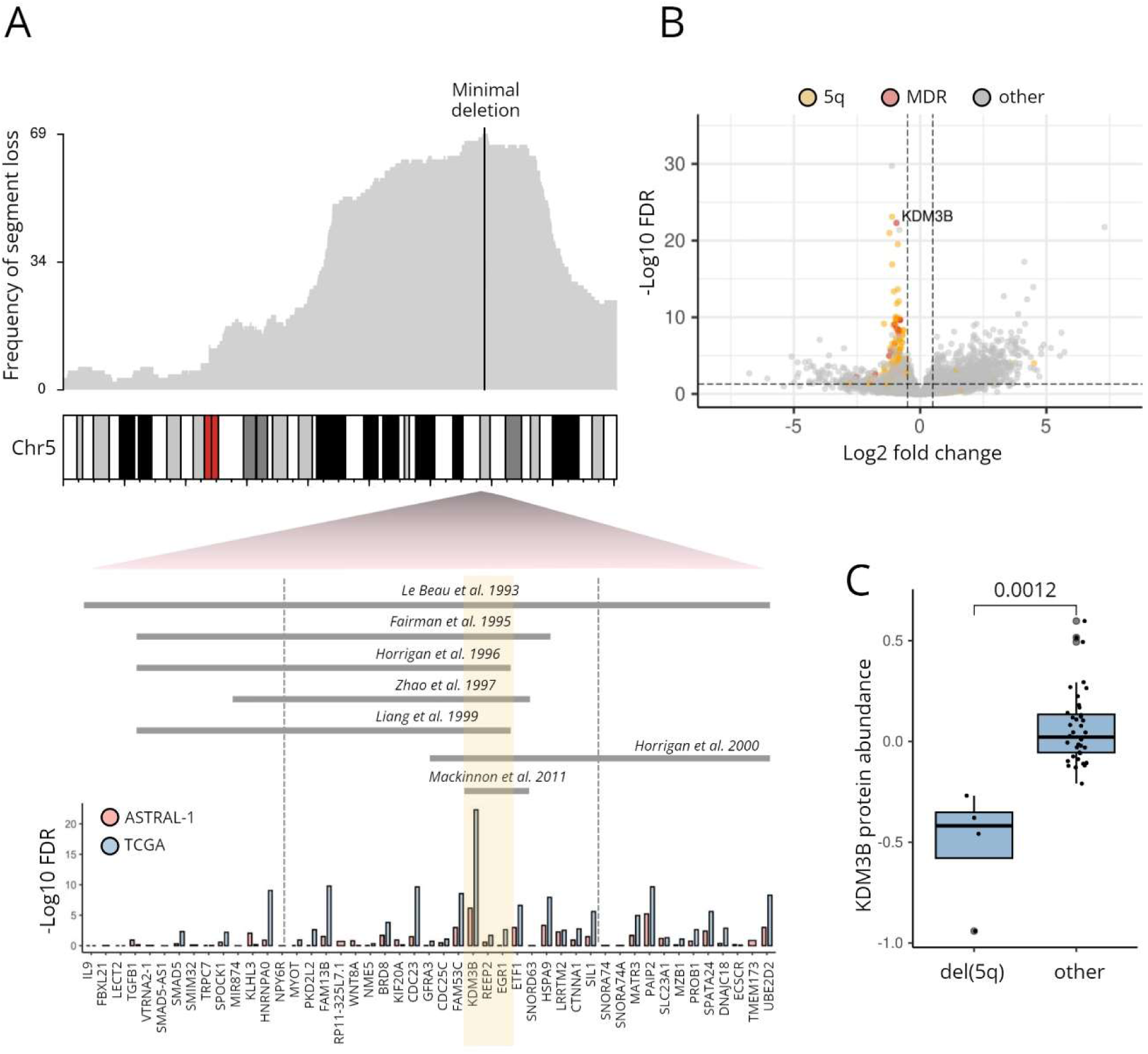
Interrogation of the minimally deleted region in del(5q) AML implicates KDM3B as a likely haploinsufficiency candidate. **A.** Density plot showing the frequency of deleted segments on chromosome 5 among del(5q) samples in our cohort. The location of the CDR at 5q31.2 is shown. Minimally deleted intervals described in the literature are indicated in the zoomed region below. Dotted gray lines indicate the boundaries of the minimally deleted region that could be defined in our cohort. Barplots show the -log10 adjusted p-value from differential expression analysis comparing del(5q) to other AML in our (pink) and TCGA (blue) cohort, for all genes within the minimally deleted region. **B.** Volcano plot resulting from differential expression analysis comparing del(5q) to other AML cases in the TCGA RNA-seq dataset. Genes located on chromosome 5q (yellow) and within the minimally deleted region (red) are highlighted. **C.** Boxplot comparing the protein abundance of KDM3B in del(5q) and other AML patient samples from Kramer et al. ^49^.

Since KDM3B is also the only epigenetic regulator encoded from the CDR, we reasoned that the loss or haploinsufficiency of this gene might explain the observed methylation signature in del(5q) patients. Specifically, we hypothesized that depletion of KDM3B should result in increased H3K9me1/2, and consequently trigger *de novo* DNA methylation at its targeted regions. In line with this we also found a significant enrichment of genes within lamina-associated domains (LADs; regions associated with H3K9me2) among LMC3-hypermethylated genes as well as among genes downregulated in del(5q) compared to other ckAML patients (Figure S7).

We also noted that del(5q) occurs in a mutually exclusive pattern with mutations in IDH1/2 (Figure S8A), while these two subgroups share partially overlapping patterns of DNA hypermethylation, which can be seen upon clustering of AML bulk methylomes (Figure S8B). These observations might reflect the metabolic dependency of KDM3B on alpha-ketoglutarate ^5,52,53^, similar to what has been described for the mutually exclusive TET2/IDH mutations ^5^. Since alpha-ketoglutarate is depleted in IDH mutant tumors, the activity of KDM3B and other histone lysine demethylases would be therein impaired. In this way, del(5q) and mutIDH may converge on partially overlapping epigenetic disruptions.

Collectively, these data lead us to argue that *KDM3B* is the most plausible target of the 5q deletion, and to reason that its depletion might explain the aberrant DNA methylation pattern in this subgroup.

### The del(5q) methylation signature correlates with expression of other H3K9me1/2 regulators and the *de novo* DNA methyltransferase *DNMT3B*

Beyond the effects of KDM3B depletion, we speculated that the variability in LMC3 levels within and outside of del(5q) patients might be influenced by differences in the expression of related epigenetic regulators. To explore this, we compiled a list of genes known to participate in epigenetic processes and tested each gene’s expression for a correlation with LMC3 (Figure 4A, Table S17). Across multiple datasets, *KDM3B* was the epigenetic regulator exhibiting the strongest negative correlation. Among the strongest positive correlations, we noticed the *de novo* DNA methyltransferase, *DNMT3B* (Figure 4A & Figure S9A). This was also reflected at the protein level (Figure S9B). Since DNMT3B is known to act at sites of H3K9me1/2 and to interact with the H3K9me1/2 methyltransferase G9a ^54,55,56,57^, we hypothesized that its overexpression could contribute to the del(5q) hypermethylation signature downstream of KDM3B depletion (Figure 4B).

**Figure 4.**
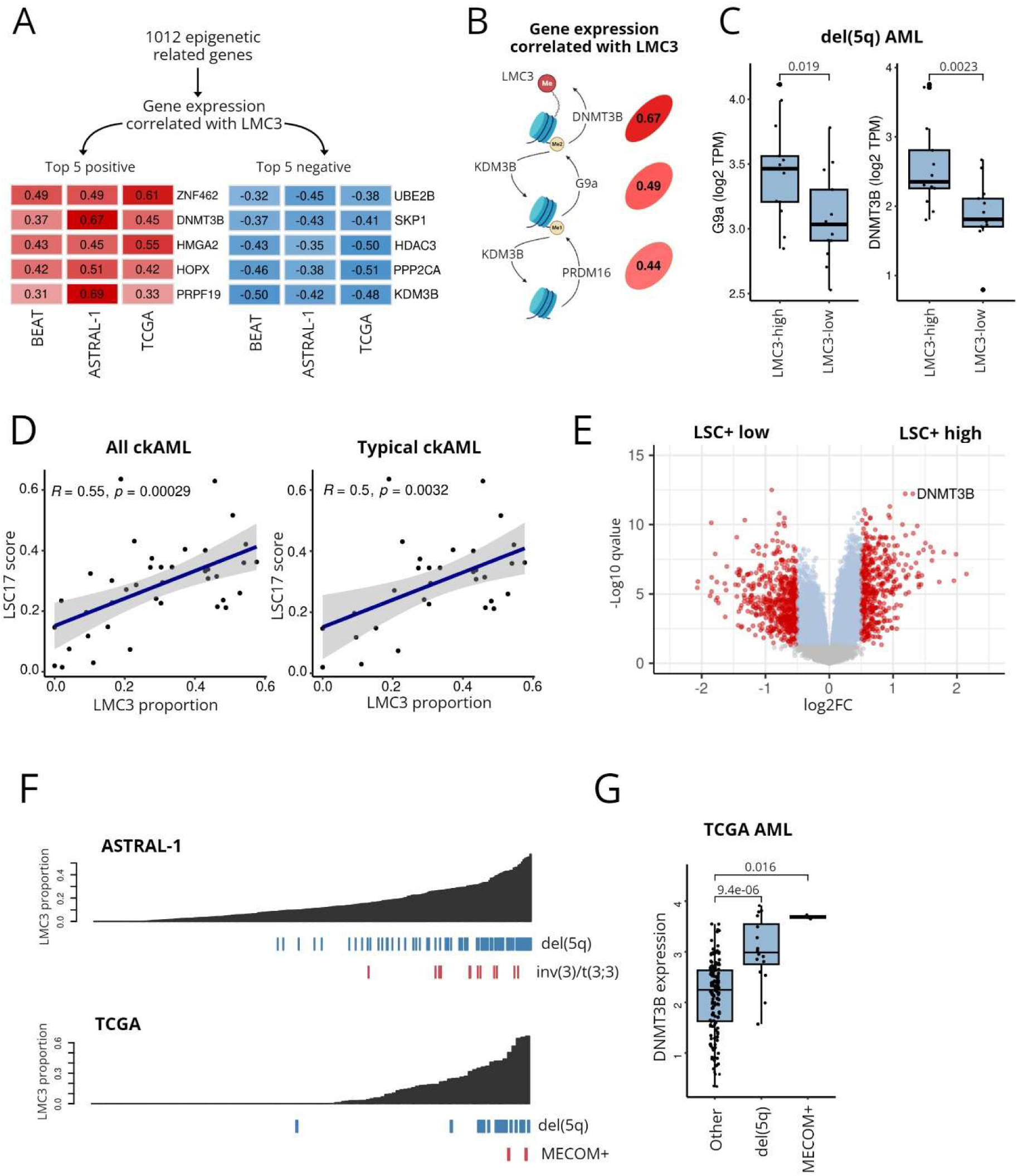
The del(5q) methylation signature correlates with expression of other H3K9me1/2 regulators and the downstream effector of de novo DNA methylation, DNMT3B. **A.** Gene expression of all epigenetic-related genes was tested for correlation with LMC3 proportion in our (ckAML), TCGA and BEAT AML cohorts. Shown are the Pearson correlation coefficients for the top 5 positive and negative correlations, ranked by mean correlation across the three cohorts. **B.** Correlation plot depicting Pearson coefficients for the correlation of DNMT3B, G9a and PRDM16 with LMC3 proportion in our ckAML cohort. **C.** Boxplots comparing the gene expression (log2 transformed TPM) of G9a and DNMT3B in del(5q) samples separated by median LMC3 proportion. Wilcoxon’s p-values shown. **D.** Scatter plots showing the correlation between LMC3 proportion and LSC17 score among ckAML (left) and among typical ckAML (right) from our cohort. Pearson coefficients and p-values are indicated. **E.** Volcano plot resulting from differential expression analysis of LSC+ vs LSC-AML cells using gene expression data from the Ng *et al.* LSC17 study. **F.** Barplots showing association of LMC3 with MECOM overexpression and the associated t(3;3)/inv(3) in our own and TCGA AML cohorts. Samples are ordered by increasing LMC3 proportion. **G.** Boxplot comparing gene expression of *DNMT3B* (log transformed RPKM) in del(5q), MECOM-high AML and other AML samples from the TCGA AML cohort.

To further support this model, we found that LMC3 correlated with increased expression of two H3K9me1/2 methyltransferases; *EHMT2* (G9a) and *PRDM16* (Figure 4B), both of which can oppose the activity of KDM3B. We also estimated LMC proportions in an additional AML sample in which the H3K9me1 methyltransferase PRDM16 was overexpressed to levels higher than all other AML samples in our ckAML cohort (List *et al.,* manuscript under review). Here we found LMC3 levels comparable to that in del(5q) AML (Figure S9C), further suggesting that LMC3 reflects an increase in H3K9me1/2. *DNMT3B* and *EHMT2* expression also maintained a correlation with LMC3 independently of del(5q) status, i.e. del(5q) patients with above-median LMC3 proportion had significantly higher expression of both *DNMT3B* and *EHMT2* compared to del(5q) patients with below-median LMC3 proportion (Figure 4C).

Highlighting the significance of *DNMT3B* in AML, a pan-cancer TCGA analysis revealed higher *DNMT3B* expression in AML than in any other tumor type, except tumors of embryonic origin (Figure S9D). Moreover, *DNMT3B* is among the 17 genes that contribute to the prognostic stemness signature, LSC17, as well as other previously established stemness signatures in pediatric AML ^48,58^, and in line with this, we noted a strong correlation between LMC3 and LSC17 (Figure 4D & Figure S9E). Since others have pointed out high LSC17 as a feature of “typical” ckAML (ckAML encompassing del(5q)/(7q)/(17p))^59^, we repeated this analysis on the subgroup of typical ckAML and found that LMC3 proportion correlated with LSC17 independently of the typical/atypical classification, i.e. LSC17 is higher in del(5q) patients compared to typical ckAMLs in which 5q is retained (Figure 4D). Reanalysis of gene expression data from the LSC17 study ^48^ revealed *DNMT3B* as the single most significantly upregulated gene in LSCs compared to their leukemic blast counterparts (Figure 4E, Table S18).

To understand the mechanisms leading to *DNMT3B* overexpression, we inspected its methylation status. We found that hypomethylation of a CpG site within the *DNMT3B* promoter in del(5q) patients correlated with its gene expression, suggesting that *DNMT3B* itself might be activated through epigenetic mechanisms in this subgroup (Figure S9F). We examined the methylation status of this region in methylomes from sorted LSCs and blasts from a del(5q) patient ^38^. Only LSC fractions were hypomethylated at this region (Figure S8G), suggesting that overexpression of *DNMT3B* might specifically be a feature of del(5q) LSCs, resulting in a methylation pattern that is inherited by their blast progeny.

AML overexpressing the *MECOM/EVI-1* oncogene is an aggressive subgroup that was previously shown to exhibit a distinct hypermethylation signature ^13^, mediated by MECOM’s interactions with DNMT3B as well as H3K9 methyltransferases ^60^. To support our hypothesized role for DNMT3B in bringing about the LMC3 methylation signature, we considered whether a similar epigenetic pattern might be detected in this subgroup. Indeed we found that AML overexpressing *MECOM*, or carrying the associated t(3;3)/inv(3) alterations, carried significantly higher levels of LMC3 compared to all other subgroups without 5q deletions (Figure 4F, Figure S10A). In line with this, overexpression of *DNMT3B* was common to both del(5q) and *MECOM*-overexpressing AML (Figure 4G, Figure S10B). We deduce that del(5q) and *MECOM*-overexpressing AML likely converge on a common epigenetic mechanism involving the methyltransferase DNMT3B, giving rise to overlapping patterns of DNA hypermethylation.

Overall these data suggest that DNMT3B may be an important player in del(5q) AML, likely contributing to the LMC3 methylation signature, and possibly also to stemness features of the del(5q) phenotype.

## Discussion

Previous efforts to characterize the DNA methylation landscape of AML have succeeded in linking many of the most common mutational and cytogenetic events to distinct epigenetic patterns ^10,11,12^. Here, we focused on a large cohort of elderly AML patients^32^, which are enriched for different molecular and cytogenetic alterations, including a higher frequency of ckAML. There have been limited attempts to further categorize this heterogeneous subgroup, excepting the distinction of “typical” and “atypical” ckAML, the former defined by the loss of material on chromosome 5/7q and/or 17p, a higher frequency of TP53 mutations, and an especially dismal clinical outcome ^15,16^. Improved characterization of ckAML is needed in order to untangle the molecular events linking recurrent chromosomal aberrations to profound genomic instability and chemoresistance, and ultimately to point towards novel therapeutic avenues in this domain. While ckAML has been linked to DNA methylation patterns before ^11,12^, it has never been comprehensively characterized from an epigenetic perspective.

The mechanisms underlying some of the most recurrent CNAs in AML are still not understood. Among these, del(5q) is the most common, where it occurs in 60-80% of ckAML, and occasionally as a sole abnormality ^15^. Despite the widespread disruption of epigenetic enzymes in almost all other AML subgroups, an epigenetic basis for del(5q) has never been considered. Here, we discovered that del(5q) AML is an epigenetically distinct subgroup defined by a unique signature of DNA hypermethylation. Reconsidering this subgroup from an epigenetic perspective, we find multiple lines of evidence suggesting that the H3K9me1/2 demethylase *KDM3B,* is the most plausible del(5q) target.

A few previous studies have also drawn attention to KDM3B in AML and highlighted its importance in hematopoietic development. Most notably, evidence for tumor suppressor activity of KDM3B in del(5q) cell lines was suggested in a 2018 study by Xu *et al.*^61^. Nevertheless, *KDM3B* has received very little attention in studies of del(5q) AML relative to other CDR genes such as *EGR1* and *CTNNA1* ^61^. More recently, Waarts *et al.* identified KDM3B as a selective dependency of IDH/TET-mutant HSCs ^62^, which is intriguing considering the mutually exclusivity of del(5q) and IDH mutations which we observed. A role for KDM3B in regulating MLL/KMT2A rearrangements in AML was also recently highlighted by Gray *et al.* ^63^, and *KDM3B* depletion was also shown to promote genomic instability in other cancers, which is interesting considering the selection for del(5q) in the context of a complex karyotype ^64^. Other regulators of H3K9me1/2 are known to be disrupted in AML; for example, overexpression of the methyltransferases PRDM16 and G9a, has been linked with poor outcome and features of leukemic stemness ^65,66,59^. Outside of malignancy, patterns of H3K9me1/2 have been shown to regulate lineage commitment in hematopoietic stem cells^67^, and KDM3B itself was shown to be important in regulating hematopoietic development through its effect on histone lysine and arginine methylation ^68^. This background strengthens the argument that KDM3B disruption, even to the degree of haploinsufficiency, could have profound phenotypic effects during hematopoietic development. Although, given the typically large size of 5q deletions, it is difficult to rule out cooperative or combined effects of multiple haploinsufficient genes, our findings highlight the need for follow-up studies to disentangle the role of H3K9me1/2 in AML pathogenesis.

We have also proposed that the methylation signature in del(5q) AML is likely mediated by the *de novo* DNA methyltransferase DNMT3B, which results in a pattern of methylation resembling that of *MECOM/PRDM16*-overexpressing AML. DNMT3B is predominantly active during embryogenesis, but is also aberrantly expressed in some tumor types ^48,69^ and is emerging as a potentially important player in AML, where its expression is higher than other non-embryonic tumors. While *DNMT3B* has been linked to features of leukemic stemness and poor prognosis ^48,7071^, we are the first to identify a DNA methylation signature associated with DNMT3B in AML, and to link DNMT3B to the del(5q) subgroup. While interactions between DNMT3B and H3K9me1/2 have been extensively documented ^54,55,56,57^, addressing the mechanistic link between KDM3B depletion and DNMT3B overexpression will be an important future goal.

As well as identifying the del(5q) methylation signature, our DNA methylation analysis also represents the first comprehensive epigenetic characterization of AML in the elderly. This resulted in a number of interesting observations; for example, co-occurring mutations in *DNMT3A/IDH* or *DNMT3A/TET2* tended to result in methylation signatures resembling normal HSPCs. While a similar so-called “epigenetic antagonism” was previously described by Glass et al. for *IDH,* we show that this phenomenon extends also to *TET2* mutations ^11^. Our analysis also highlighted the previously unrecognized epigenetic similarity of AML with aberrations affecting the RUNX1 signaling network, including *RUNX1* and *CBFβ* mutations, del(21q), and inv(16).

## Conclusion

We have shown that del(5q) AML is an epigenetically distinct subgroup, defined by a unique DNA methylation signature, which we link to reduced activity of the H3K9me1/2 demethylase KDM3B, and *DNMT3B* overexpression. Our findings shed new light on a highly aggressive and poorly understood AML subgroup and highlight a need to further explore epigenetic mechanisms of leukemic progression in ckAML.

## Materials & Methods

### DNA methylation analyses Methylation EPIC array preprocessing

DNA methylation profiles were generated using the Infinium MethylationEPIC array for 477 AML samples from the ASTRAL-1 cohort (described by Jahn et al.^7^). Quality control, preprocessing and normalization were performed using RnBeads 2.0 ^72^. Filtering was applied to remove SNP-overlapping probes, cross-reactive probes, sites outside of CpG context and CpGs mapping to sex chromosomes, as well as probes with detection p-value > 0.05 and sites covered by fewer than three beads. Beta-values were normalized and background subtraction was carried out using the “scaling.internal” and “sesame.noobsb” methods, respectively.

### Methylome deconvolution

Methylome deconvolution was performed using MeDeCom^31^ according to an established protocol^33^. MeDeCom is a reference-free deconvolution method based on constrained non-negative matrix factorization, which aims to decompose of a set of bulk methylomes to recover their constituent latent methylation components (LMCs) and the proportions of LMCs within each patient sample (Figure 1A). For deconvolution, the 20,000 most variably methylated CpG sites across all samples were selected, in order to limit computational demands while retaining the most informative features in the dataset. Potential confounding factors including age, sex, batch, Sentrix ID and Sentrix position were adjusted for using Independent Component Analysis ^73^. Deconvolution was performed using a range of K values from 2 to 15. K = 11 was selected as the optimum number of LMCs based on the cross-validation error (Figure S1A). The regularization parameter lambda = 0.01 was selected in order to minimize the cross-validation error (Figure S1B).

### Estimating sample purity

Sample purity was estimated from the methylation data using the InfiniumPurify method ^74^. InfiniumPurify infers tumor purity in a cancer-type specific manner based on a kernel density estimation method, using a predefined set of CpG sites differentially methylated between tumor (AML) and corresponding normal tissue. Samples with low estimated purity (lower tertile) were excluded from downstream analyses to avoid spurious interpretations.

### Prediction of Leukemic Cell of Origin

A predicted cell of origin was assigned to each AML sample according to previously established methods ^38^. HumanMethylation450K data from sorted hematopoietic progenitor cell states from healthy donors, including hematopoietic stem cells (HSCs), multipotent progenitors (MPPs), lymphoid-primed multipotent progenitors (LMPPs), common myeloid progenitors (CMPs), granulocyte/monocyte progenitors (GMPs) and megakaryocyte/erythroid progenitors (MEPs), was used as a reference. A set of 216 differentially methylated regions were identified by Jung *et a*l. from pairwise comparison between the differentiation states^38^. Hierarchical clustering of AML together with normal progenitor cell samples, on the methylation of CpGs sites within these regions, resulted in three groups of GMP-like (more differentiated), MEP/CMP-like and MPP/LMPP-like (less differentiated) AMLs (Figure S1D). LMC proportions were compared between these three groups by Wilcoxon’s test.

### Estimation of LMC proportions in external datasets

To validate our interpretations of LMCs, the factorize regression function from the MeDeCom R package was used to derive an estimate of the LMC proportions in two publicly available datasets comprising HumanMethylationEPIC/450K data from the TCGA and BEAT-OSU AML cohorts. Clinical and molecular annotations for TCGA-AML were retrieved from the Genomic Data Commons using the TCGAbiolinks package, and for BEAT AML were obtained from the supplementary tables of Tyner et al. ^75^.

### Biological interpretation of LMCs

To identify LMCs derived from non-leukemic cell types, LMC proportions were correlated against methylomes of normal hematopoietic cells using EPIC array data from Salas et al. ^34^. For further interpretation, sets of LMC-specific hypomethylated and hypermethylated CpG sites were defined for each LMC as those with a methylation beta value > 0.5 above or below the mean of the remaining LMCs. Gene ontology and transcription factor binding site enrichment analyses for the resulting CpGs were performed using the clusterProfiler^76^ and LOLA^77^ R packages. To identify LMCs associated with mutations or cytogenetic subgroups, LMC proportions were compared between groups using Wilcoxon tests.

### Defining LMC-based subgroups

To define LMC-based subgroups, consensus k-means clustering ^42^ was applied to the matrix of LMC proportions, excluding the lymphoid-like (LMC9) and neutrophil-like (LMC1) components. Euclidean distance and complete inner linkage were used for clustering and k = 9 was selected as the optimum number of clusters based on the cumulative distribution function (CDF) (Figure S1G).

### Differential methylation analysis

A set of CpG sites which are differentially methylated in LMC3-high AML was defined by comparing the methylomes (n=73 samples) of the LMC3-high AML cluster (from consensus clustering, after excluding low purity samples) to (i) all remaining AML clusters (n=244 samples), and (ii) normal HSCs (n=10 samples). By this strategy we were no longer restricted to those 20,000 CpG sites used for deconvolution, and could consider methylation changes from a normal state as well as between malignant states. Differential methylation between groups was computed at the level of CpG sites using RnBeads. CpG sites with absolute mean beta value difference > 0.2 and adjusted p-value < 0.05 were considered differentially methylated. CpG sites which were consistently hypo/hypermethylated in both comparisons were taken forward for enrichment analyses. Among differentially methylated sites, enrichments of transcription factor binding sites and gene ontologies were calculated using the LOLA^77^ and clusterProfiler^76^ R packages, respectively. The list of homeobox genes was retrieved from Wilming et al.^75,78^ and tested for enrichment among LMC3 hypermethylated CpGs using the hypergeometric test.

### RNA-seq analysis

Differential gene expression analysis using RNA-seq datasets from our and TCGA cohorts was performed using DEseq2 with default parameters^79^. Pan-cancer and pan-tissue gene expression analyses for *KDM3B* and *DNMT3B* were performed using TCGA and GTEx data from the GEPIA portal ^80^. To investigate pathway-level gene expression changes associated with the del(5q) methylation signature, gene set variation analysis (GSVA) was performed using the GSVA R package^81^ and the resulting pathway-level expression scores were tested for correlation with LMC3 proportion. Stemness scores were computed for each patient using the weighted sum of expression of the 17 LSC17 genes as defined by Ng *et al.* ^48^.

### Comparisons of LSCs & blasts

To compare gene expression in LSC+ and LSC- cell fractions (determined by Ng et al. based on the cells’ ability to generate an AML graft in one or more mice), microarray data were downloaded from Ng *et al.*^48^ (GSE76009) and differential expression was computed using Limma. The same dataset was used for comparisons of *KDM3B* gene expression in LSC (LSC+) and blast (LSC-) counterparts of del(5q) samples. Del(5q) status for the samples was obtained from the corresponding authors. To compare LMC3 levels in del(5q) LSCs vs blasts, 450K data from sorted LSCs (CD34+CD38+/CD38-) and paired blasts (CD34-) was downloaded from Jung et al. (GSE63409) ^38^. Conumee copy number profiles were generated to identify samples with 5q deletions ^45^. Estimated LMC proportions were derived using MeDeCom’s factor regression approach.

### Mutual exclusivity of mutations & CNV patterns

Mutual exclusivity of del(5q) and IDH mutations was determined based on the Poisson-Binomial distribution as implemented in the Rediscover R package ^82^.

### Copy number analysis and investigation of the 5q CDR

Copy number profiles were generated from EPIC array data using the Conumee R package using 20 AML samples of known normal karyotype as “flat genome” controls. The CDR was defined from our AML cohort using the Conumee segmentation results. In each sample, segments mapping to chromosome 5q were selected which had a p-value < 0.05 for loss at that region. The frequency of loss in 10kb bins was calculated and the minimal genomic interval having the highest frequency of loss was determined. This definition was compared to that of six studies of del(5q) AML or high-risk MDS identified from a literature search, which each proposed a variation on the minimal interval of interest, largely overlapping with our own definition. For all downstream analysis of CDR genes, the 50 genes flanked by *Il9* and *UBE2D2* were considered as an accepted and conservative definition of the CDR ^83,23^. The expression of CDR genes was correlated against LMC3 proportions using Pearson correlation.

### Protein expression analysis

Protein expression of CDR genes was examined using proteome data from the Cancer Cell Line Encyclopedia (CCLE) ^51^. AML cell lines were selected using the DepMap portal and identified as del(5q) or otherwise. Protein expression was compared between groups by Wilcoxon test, excluding outliers from the analysis if their expression was > 3 standard deviation from the mean of all AML cell lines. Tandem mass tag (TMT) proteome data for 44 AML patients was retrieved from Kramer *et al.*^49^ Abundance values were calculated from reporter-ion intensity, log2 transformed and median centered at 0. A second mass-spectrometry-based proteome dataset of 177 AML samples including 3 del(5q) samples was obtained from Jajavelu et al. ^50^.

### Survival and drug sensitivity analyses

Clinical data from the BEAT AML cohort were used to assess the prognostic value of LMCs. Hazard ratios and p-values were computed for each LMC using cox regression. To further assess the prognostic significance of LMC3, samples were separated into LMC3-high (above mean) and LMC3-low (below mean) groups, and the LMC3-high group was further subdivided into del(5q)- positive and del(5q)-negative patients. Conumee copy number profiles were used to reliably define samples as del(5q) or otherwise. Pairwise Log rank test p-values were computed using the survival and survminer R packages.

Ex vivo drug sensitivity data for 122 small molecule inhibitors was obtained from the BEAT AML cohort ^75^. To compare global drug sensitivities between patient subgroups, binary sensitive or resistant calls were assigned for each sample to each drug, considering samples with the highest 20% of AUC values for a given drug as sensitive. Then for each sample the proportion of drug sensitivities was quantified, and these proportions were compared between LMC3-high and LMC3-low subgroups.

### Data and code availability

DNA methylation data for the 477 AML samples and scripts used throughout the manuscript will be made publicly available upon publication of the manuscript. All publicly available datasets used in the study are listed in the supplementary tables (Table S1).

## Acknowledgements

We thank the Genomics and Proteomics Core Facility, the Omics IT and Data Management Core Facility, and the Microarray Core Facility of the DKFZ Heidelberg. We thank Dieter Weichenhan for comments on the manuscript, Jean Wang and Amanda Mitchell for providing cytogenetic annotations for samples from the Ng *et al.* dataset. This work was supported in part by the German Funding Agency (DFG) through FOR2674 (to CP) and SFB 1074 (to HD, KD, CP), the Carreras Foundation (CP), and the Helmholtz Foundation. MS is supported by a postdoctoral fellowship by the Dr. Rurainski Foundation. Biobanking of patient samples from the ASTRAL-1 trial was generously supported by a grant from ASTEX Pharmaceuticals.

