## Supplementary Figures for "Acute Myeloid Leukemia with deletion 5q is an epigenetically distinct subgroup defined by heterozygous loss of *KDM3B*"

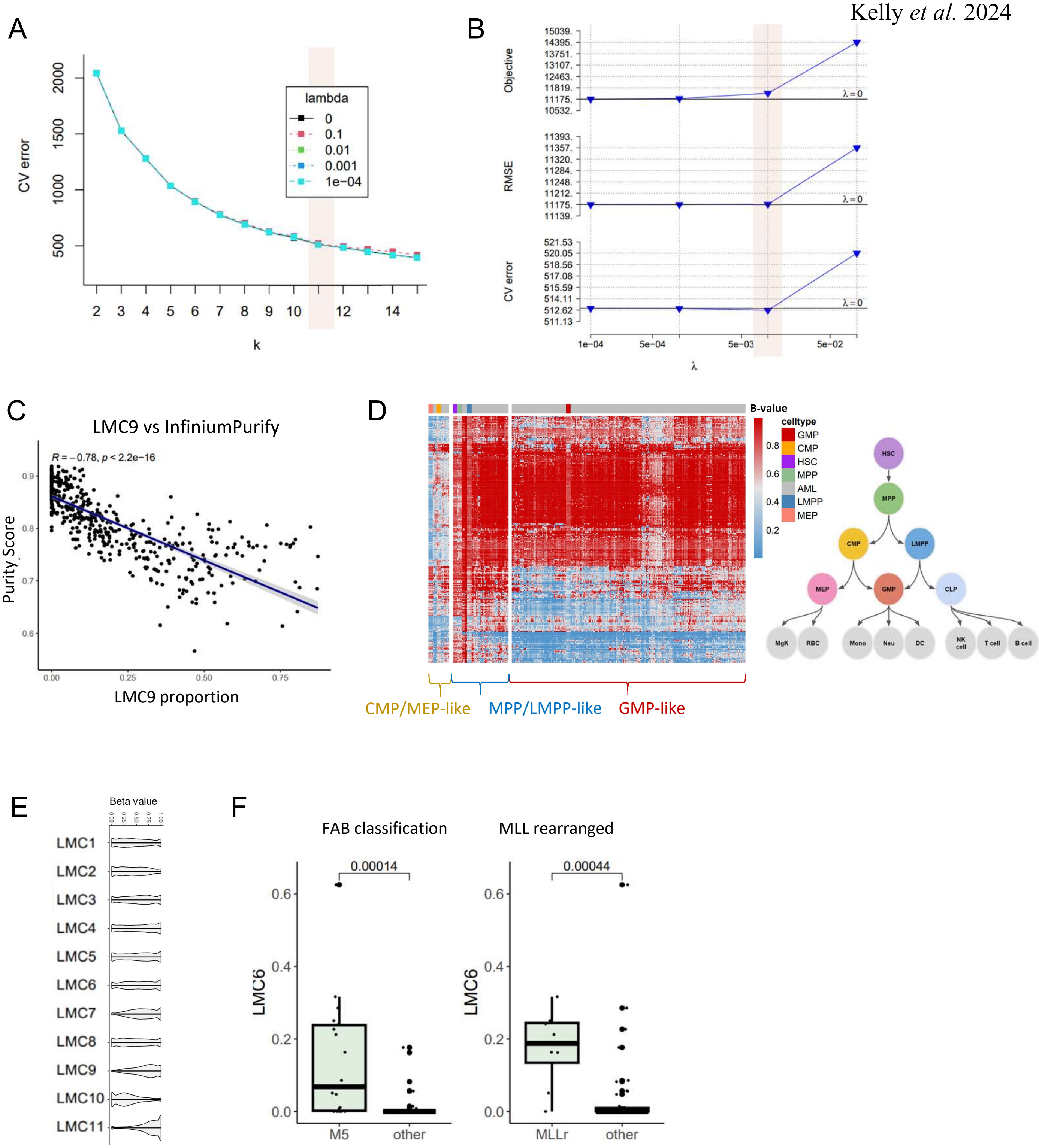


**Figure S1. Methylome deconvolution-based epigenetic characterisation of Acute Myeloid Leukemia**
**A.** Plot of cross validation (CV) error used for K selection over 2-15 LMCs. The selected K value is highlighted. **B.** Selection of regularisation parameter Lambda (log scale) over a range of 0 to 0.1. The selected Lambda value is highlighted. **C.** Scatter plot correlating LMC9 proportion with sample purity estimated by InfiniumPurify. Pearson correlation coefficient and p-value shown. **D.** Heatmap obtained from hierarchical clustering of AML methylomes together with hematopoietic stem cell (HSC), multipotent progenitor (MPP), lymphoid-primed multipotent progenitors (LMPP), common myeloid progenitors (CMP), megakaryocyte/erythroid progenitors (MEP) and granulocyte/monocyte progenitors (GMP). Samples are clustered based on 216 regions of differential methylation between progenitor cell states. The resulting GMP-like, MPP/LMPP-like and CMP/MEP-like clusters serve as an estimate of the AML cells of orgin. **E.** Violin plots comparing the global methylation levels of each LMC based on the beta values of all 20,000 CpG sites used for methylome deconvolution. **F.** Boxplots comparing LMC6 proportion in M5 vs other FAB cateogry AML and MLL/KMT2A rearranged vs other AML in the BEAT-OSU AML cohort. Wilcoxon's p-value shown.

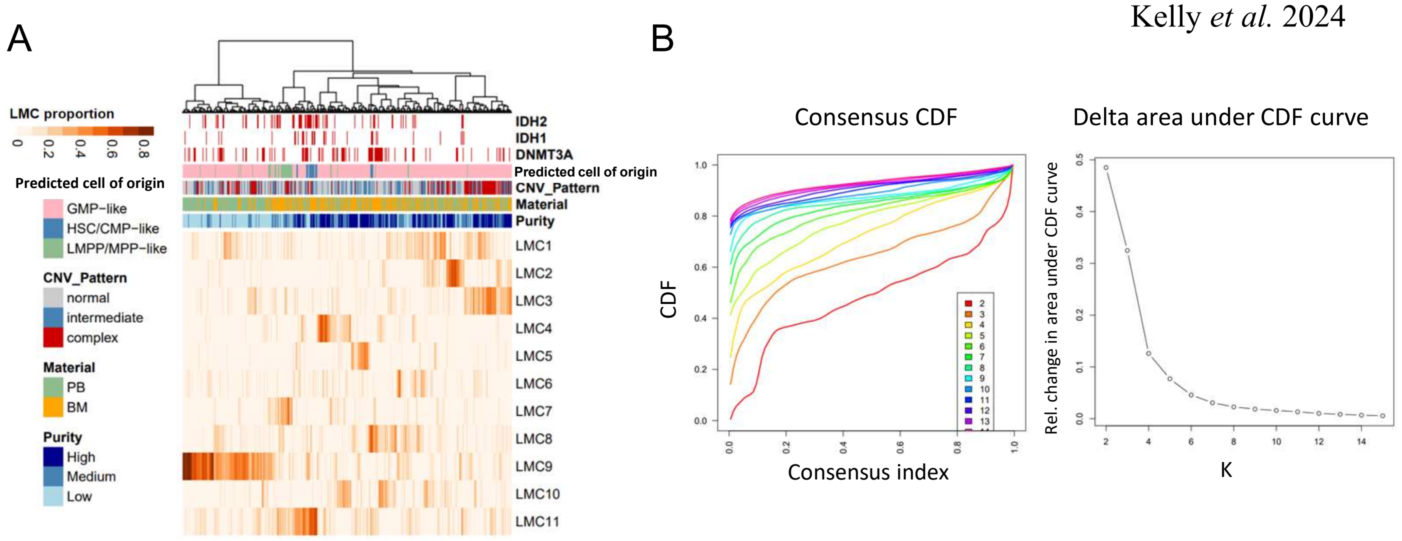


**Figure S2. Clustering of AML patient samples based on methylome deconvolution**

**A.** Heatmap of LMC proportions for the 11 LMCs defined in ASTRAL-1 AML cohort, before removal of LMC1 and LMC9 for consensus clustering. Hierarchical clustering was performed using the ward.D2 method. PB=peripheral blood, BM=bone marrow. High, low, and medium purity represents InfiniumPurify scores separated into tertiles. **B.** Plots of the cumulative distribution function (CDF) and area under the CDF curve, used for K selection in consensus clustering.

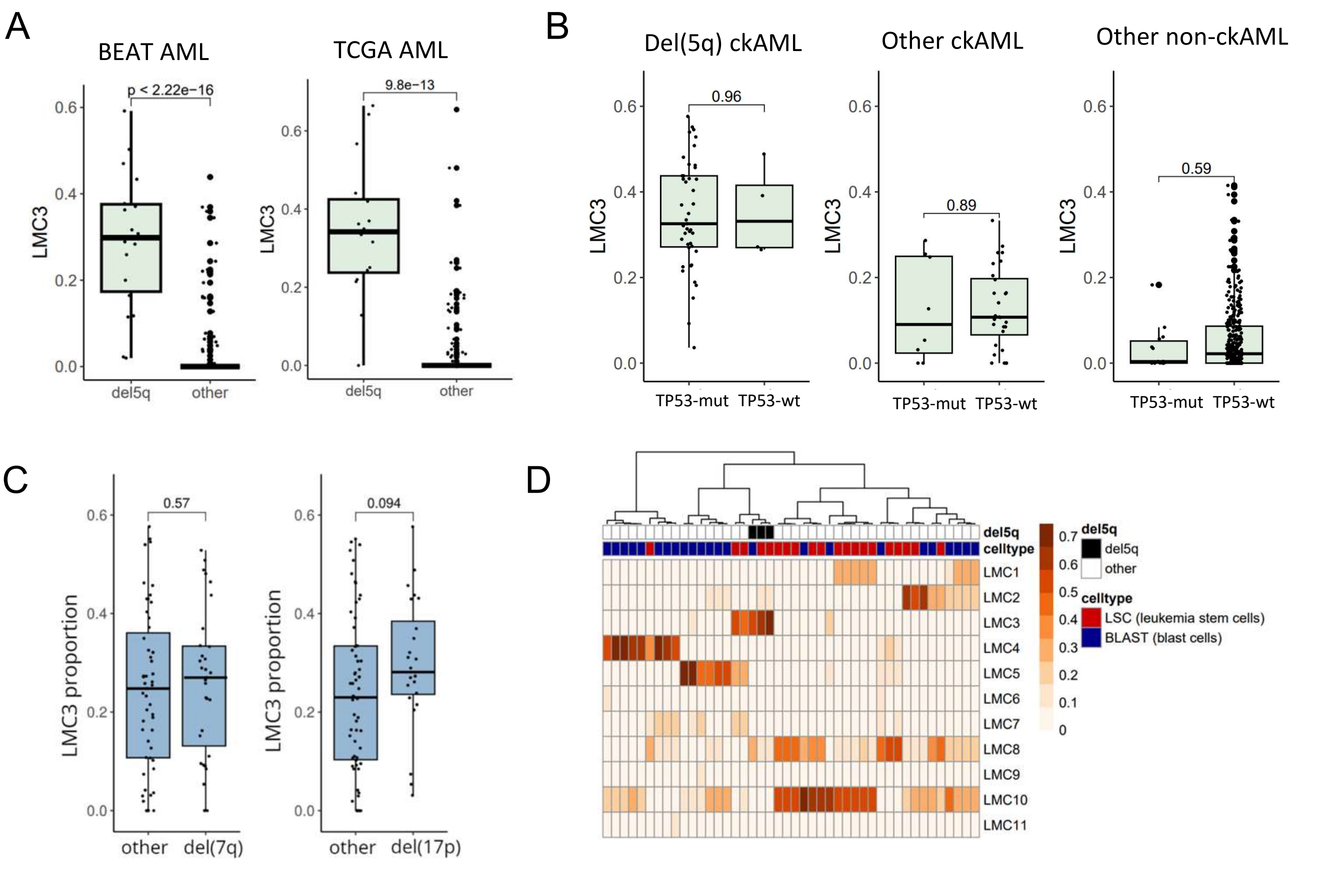


**Figure S3. Del(5q) AML is defined by a unique DNA methylation signature**
**A.** Boxplots comparing LMC3 proportion in del(5q) and 5q-retained AML samples from BEAT-OSU and TCGA AML cohorts. Wilcoxon's p-values shown. **B.** Boxplots comparing LMC3 proportion in TP53 mutated and wildtype AML, among del(5q) ckAML cases (left), non-del(5q) ckAML cases (middle), and non-del(5q), non-ckAML cases (right). Wilcoxon's p-values shown. **C.** Boxplots comparing LMC3 proportion among ckAML patients from the ASTRAL-1 cohort, separated according to the presence of 7q and 17p deletions. Wilcoxon's p-values shown. **D.** Heatmap showing estimated LMC proportions in sorted AML blasts and leukemic stem cells (LSCs) using 450K data from the Jung *et al.* GSE63409 dataset. Samples deriving from a del(5q) patient are annotated.

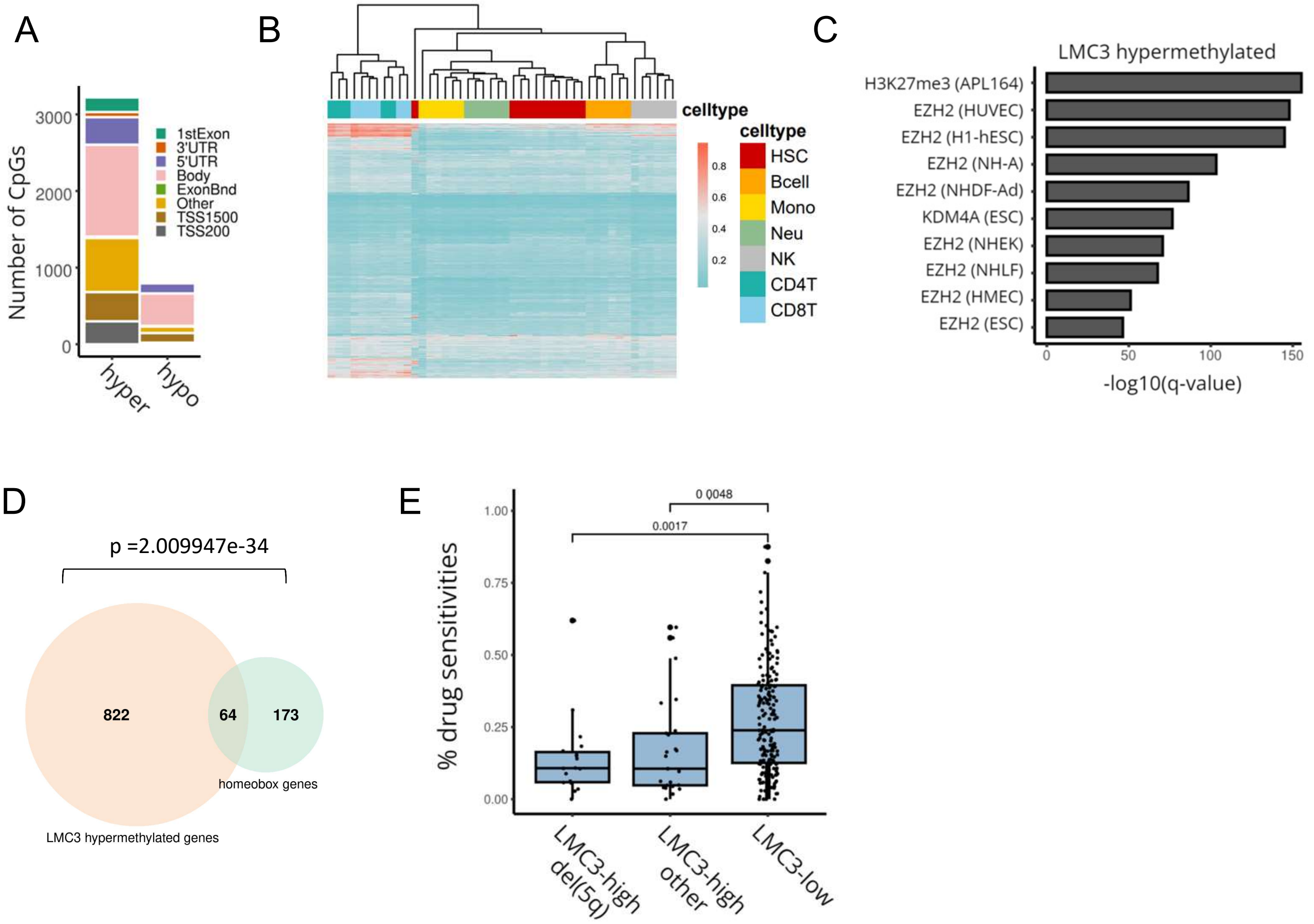


**Figure S4. Characterisation of the del(5q) methylation signature**

**A.** Stacked barplot showing the distribution of LMC3 hypo/hypermethylated CpG sites relative to UCSC RefGene group. **B.** Heatmap showing the methylation levels of LMC3-hypermethylated CpG sites in the normal hematopoietic lineage. **C.** Barplot showing top 10 locus enrichments among LMC3-high hypermethylated CpG sites. **D.** Venn diagram showing the intersection of homeobox genes among del(5q) hypermethylated CpG sites. Hypergeometric p-value is shown indicating enrichment of the gene set. **E.** Boxplot comparing overall drug sensitivity based on ex vivo drug sensitivity screens on BEAT AML samples. Patients are separated into LMC3-high and LMC3-low groups based on mean LMC3 proportion, and LMC3-high patients further stratified as del(5q) or otherwise. For each patient we quantified the proportion of drug sensitivities after assigning binary sensitive/resistant calls for each sample to each drug. Samples within the top 20% of AUC values for a given drug were considered sensitive.

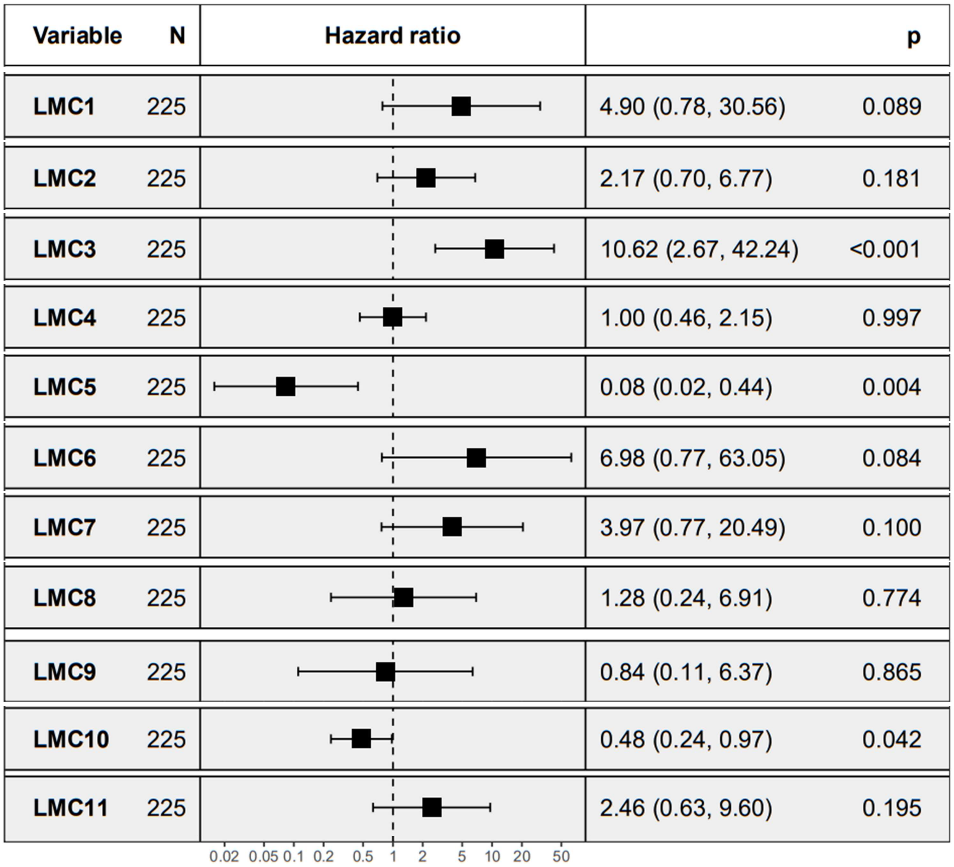


**Figure S5. Survival analysis of LMCs**
Forest plot showing the association of each latent methylation component (LMC) with overall survival using clinical data from the BEAT AML cohort. Cox proportional hazard ratio’s and p-values are shown.

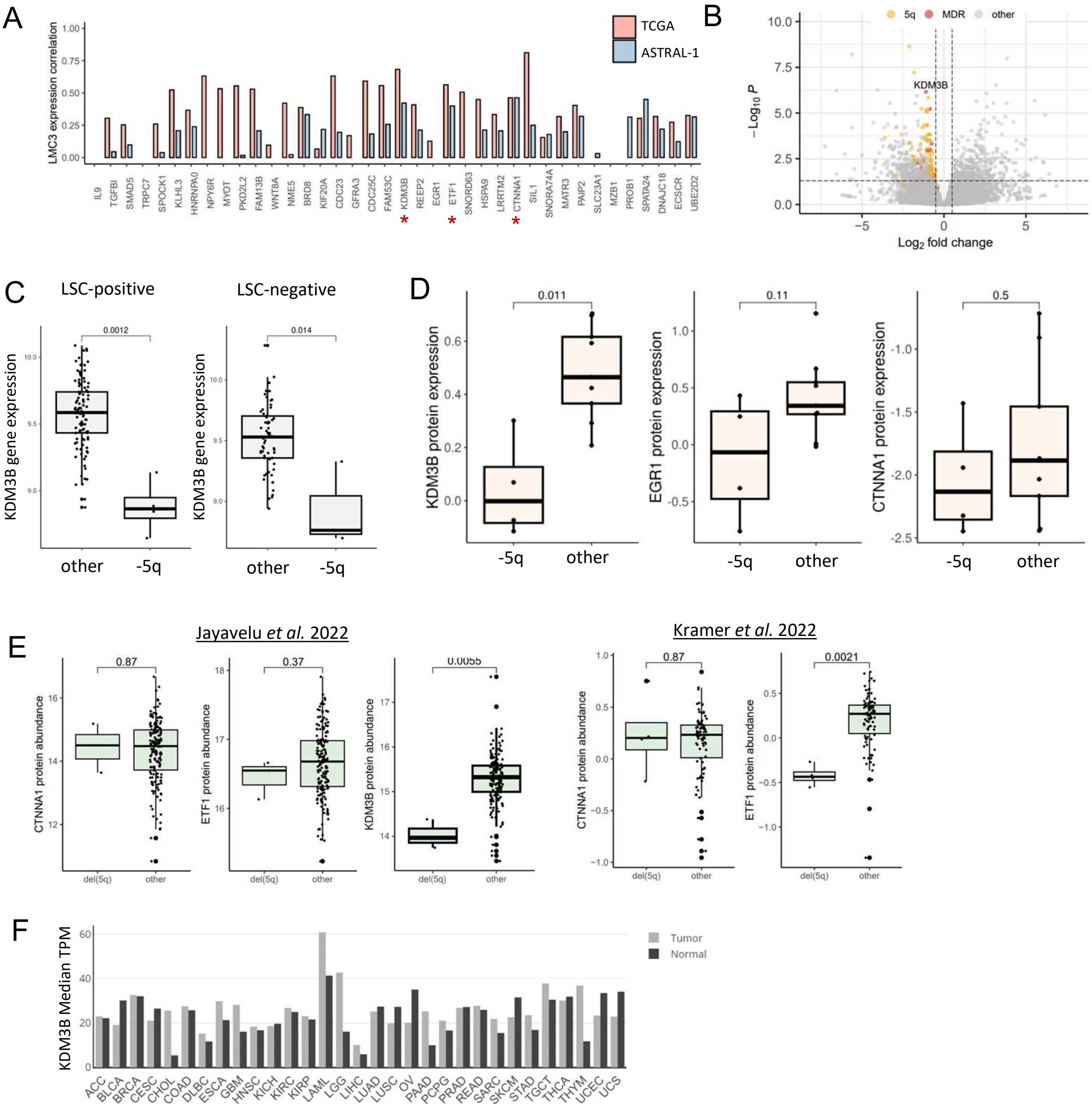


**Figure S6. The H3K9me1/2 demethyase KDM3B is a likely target of the 5q deletion**

**A.** Expression of each gene in the del(5q) commonly deleted region (CDR) was tested for correlation with LMC3 proportion in the ASTRAL-1 ckAML and TCGA cohorts. Bar plot shows the Pearson correlation coefficiencts from this analysis. The interval flanked by *IL9* and *UBE2D2* is considered as a conservative estimate of the CDR. Asterisks mark genes for which a significant p-value was detected in both datasets. **B.** Volcano plot resulting from differential gene expression analysis (DEseq2) comparing del(5q) to other ckAMLs in the ASTRAL-1 cohort. Genes located on chromosome 5q are marked in yellow and genes within the CDR described in A. are colored red. **C.** Gene expression data for sorted leukemic stem cell (LSC) positive and LSC negative AML cells from the Ng *et al.* LSC17 study were used to compare *KDM3B* gene expression in del(5q) vs 5q-retained LSC+/- fractions. Wilcoxon's p-values shown. **D.** Boxplots comparing the protein expression of candidate CDR genes, KDM3B, EGR1 and CTNNA1 in del(5q) and 5q-retained cells lines from the Cancer Cell Line Encyclopedia. Wilcoxon's p-values shown. ETF1 was not covered in these datasets so could not be included. **E.** Boxplots comparing protein expression of del(5q) candidate genes in del(5q) and other AML samples in two proteomics datasets from AML patients. Wilcoxon's p-values shown. EGR1 was not covered in these datasets so could not be included. **F.** Barplot comparing *KDM3B* gene expression across different tumor and normal tissues from the TCGA and GTEx datasets.

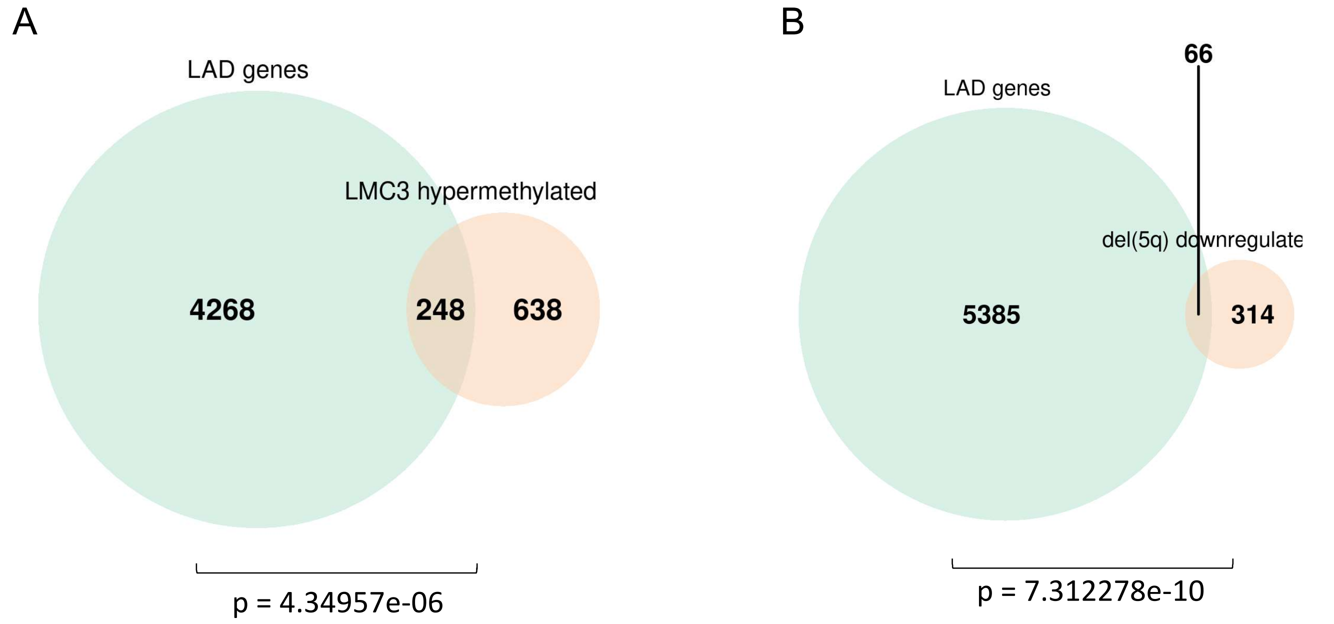


**Figure S7. Del(5q) dysregulated genes are enriched for genes within Lamina-associated domains**
**A.** Venn diagram showing the overlap of genes within lamina-associated domains (LAD genes) and genes with CpG sites hypermethylated in the LMC3-high AML subgroup. Hypergeometric p-value is shown. **B.** Venn diagram showing the overlap of genes within lamina-associated domains (LAD genes) and genes downregulated in del(5q) AML by comparison to other ckAML patients in the ASTRAL-1 cohort. Hypergeometric p-value is shown.

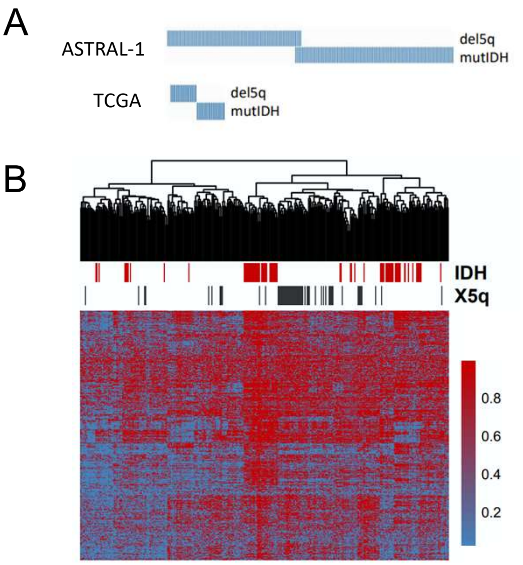


**Figure S8. Del(5q) and IDH mutation are mutually exclusive events characterised by DNA hypermethylation**
**A.** Mutual exclusivity of del(5q) and IDH (pooled IDH1/2) mutations in the TCGA and ASTRAL-1 AML cohorts. **B.** Hierarchical clustering of ASTRAL-1 AML samples based on the 5000 most variable CpG sites on the EPIC array, indicating co-clustering of del(5q) and IDH-mutated AML.

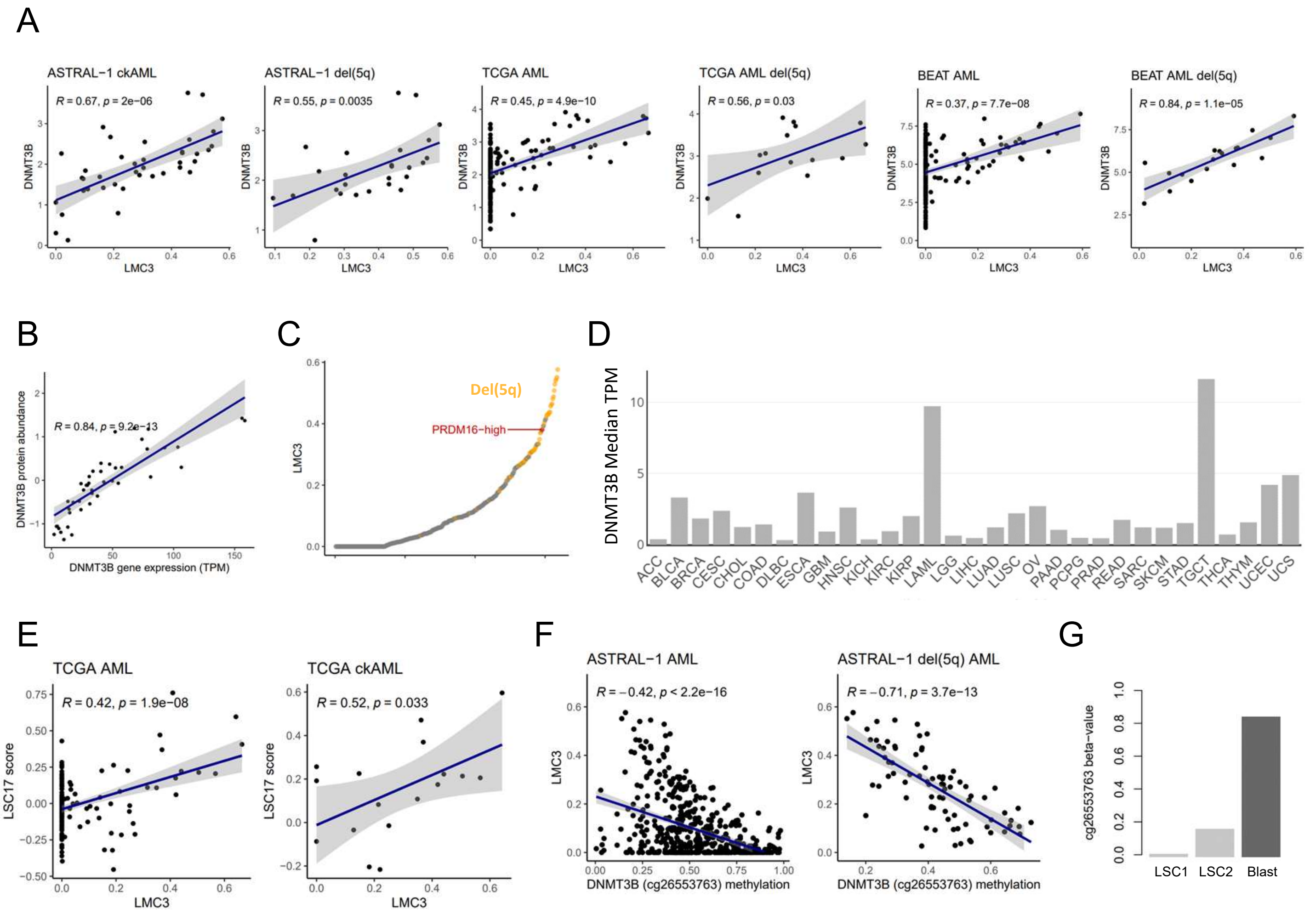


**Figure S9. DNMT3B likely contributes to the DNA hypermethylation signature in del(5q) AML**

**A.** Scatter plots correlating *DNMT3B* gene expression and LMC3 proportion in ASTRAL-1, TCGA and BEAT-OSU AML datasets, overall and among del(5q) samples. Pearson correlation coefficients and p-values are shown. **B.** Scatter plot correlating DNMT3B protein expression and gene expression across TCGA AML samples. Pearson correlation coefficient and p-value shown. **C.** Dot plot comparing LMC3 levels in del(5q) AML (yellow) and in an AML sample overexpressing H3K9me1 methyltransferase, *PRDM16* (red). **D.** Bar plot comparing *DNMT3B* gene expression (median TPM) across all TCGA tumor types. **E.** Scatter plots correlating the LSC17 score and LMC3 proportion in TCGA AML overall and among ckAML samples. Pearson correlation coefficients and p-values shown. **F.** Scatter plot correlating *DNMT3B* gene expression and promoter methylation at cg26553763 in AML samples from the ASTRAL-1 cohort. The second plot shows only those samples with del(5q). Pearson correlation coefficients and p-values shown. **G.** Bar plot comparing the methylation status of the *DNMT3B* promoter CpG site cg26553763 in leukemic stem cells (LSCs) and blasts in a del(5q) patient from the Jung *et al.* GSE63409 dataset.

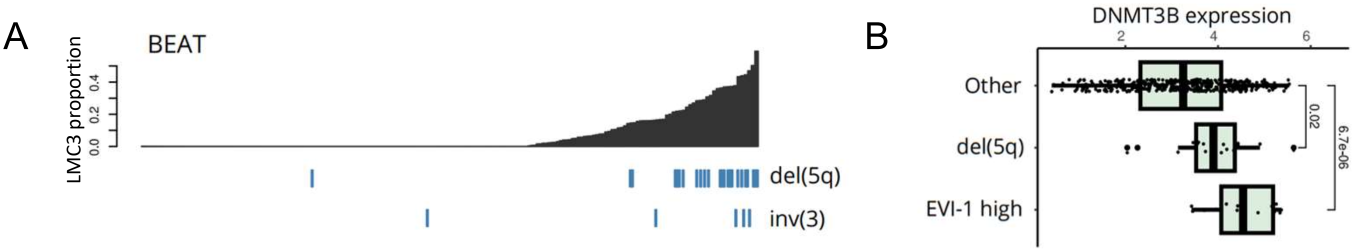


**Figure S10. Epigenetic similarities in del(5q) and MECOM+ AML converge on *DNMT3B* overexpression**
**A.** Bar plot showing association of LMC3 with inv(3) in the BEAT AML cohort. Samples are ordered by increasing LMC3 proportion. **B.** Boxplot comparing gene expression of *DNMT3B* (log transformed RPKM) in del(5q), MECOM/EVI-1-high AML and other AML samples in the BEAT AML cohort.
